# *Dlc1* controls cardio-vascular development downstream of Vegfa/Kdrl/Nrp1 signaling in the zebrafish embryo

**DOI:** 10.1101/2021.02.11.430763

**Authors:** Tanja Linnerz, Julien Y. Bertrand

## Abstract

The family of *deleted-in-liver-cancer (dlc)* genes encodes RhoGTPases and plays pivotal roles in cardiovascular development, but animal models for studying their functions are sparse due to early embryonic lethality. Gain and loss of function of *dlc1* and *dlc3* severely altered the growth of intersegmental vessels in the trunk of zebrafish embryos. Additionally, overexpression of *dlc1* affected the growth of the common cardinal veins, but could rescue the arrest of angiogenesis induced by Vegfr2 inhibition, placing *dlc1* downstream of *kdrl* signaling. Loss of *dlc1* negatively affected the lumenization of the first aortic arch arteries and the lateral dorsal aortae. *dlc1* mutants displayed a full obstruction in the early outflow tract during cardiac morphogenesis, which models to alterations in DLC1 detected in congenital heart defects in human patients. This study provides a functional *in vivo* characterization of *dlc1* and *dlc3* during vertebrate embryogenesis and places *dlc1* as a key gene to control vascular development.

## INTRODUCTION

The cardiovascular system is one of the earliest systems formed during embryonic development. Its proper formation requires a multitude of orchestrated processes, including cell proliferation, cell fate determination and migration. Many diseases develop through perturbation of these key physiological processes leading to systemic consequences. Disruptions in organ formation during embryonic development can result in congenital conditions. A well-known example is congenital heart defects (CHD), which can lead to disability or severely impaired organ function and death. Thus, studying the genetic of embryonic development is necessary to understand the mechanisms underlying these pathological conditions, thereby facilitating the development of prenatal screening tests and treatments.

The zebrafish model is suitable for studying many pathological conditions due to their conserved biology, availability of transgenic lines and optical transparency. Moreover, many genes that are embryonic lethal in mammals can be studied during early development in zebrafish larvae. One such gene, *deleted in liver cancer 1* (*dlc1*), has been shown to be a *bona fide* tumor suppressor gene and is frequently downregulated or lost in many human cancers^1-3^. The three DLC proteins, DLC1, DLC2 and DLC3, belong to the family of RhoGAP proteins and preferentially target RHOA^1^ and to a lesser degree, CDC42^1,4^. There is a high degree of similarity between all three family members, each possessing an N-terminal sterile alpha motif (SAM), an enzymatically active GAP domain and a C-terminal steriodogenic acute regulatory protein (StAR)-related lipid transfer (START) domain^5^. The function of DLC1 as a tumor suppressor is associated with its RhoGAP activity, leading to the inactivation of RHOA after concomitant binding to tensins^4^, which subsequently affects the formation of stress fibers and focal adhesions ^6^. Cytoskeletal rearrangements are key requirements in directing endothelial cell migration during blood vessel growth. Vasculogenesis and angiogenesis are thereby tightly regulated processes that depend on the interplay between numerous signaling pathways. Besides the well-known major VEGF/VEGFR signaling pathway^7^, other guidance factors, such as Semaphorins, Plexins and Neuropilins, also play a crucial role in vascular development and can additionally function as enhancing co-factors in the VEGF/VEGFR signaling axis^7,8^.

However, not much is known about the role of *dlc1* during organogenesis because its homozygous deletion is embryonic lethal in Drosophila, mouse and avian models^9-14^. *Dlc1*^*null*^ mouse embryos showed multiple anatomical changes in the developing vasculature and heart, which were analyzed postmortem due to intrauterine lethality ^9^. A recent study also linked the occurrence of a rare variant of DLC1 to human CHDs^15^. To further study the role of these genes in development, a Dlc2-KO mouse model has been established but no phenotype was detected^16^, whereas there is no current animal model for Dlc3^16^.

In this study, we performed the first *in vivo* characterization of the functional role of *dlc1* and *dlc3* (*stard8*) during embryonic and larval development using the advantages of the zebrafish model. We initially show that both genes are highly expressed in the vascular lineage. Gain and loss-of-function experiments resulted in abnormal development of the vascular network, suggesting an important role for *dlc1 and dlc3* in orchestrating endothelial cell migration during angiogenic sprouting *in vivo*. We show that *dlc1* acts directly downstream of the *kdrl* signaling axis *in vivo*, which was further confirmed by observing a loss in the integrity of the endothelial cell sheet and an uneven leading edge during common cardinal vein growth, a phenotype reminiscent of dampened Sema3/PlxnD1 signaling and RhoA inactivation ^17^. Finally, we used CRISPR/Cas9-mediated genome editing to establish *dlc1* and *dlc3* mutants. *dlc1*^null^ embryos showed defects in the lumenization of the first aortic arch arteries, in the formation of the lateral dorsal aortae and the development of the immature outflow tract in the embryonic heart. These observations are concordant with a study that recently linked DLC1 mutations to human CHDs, therefore reinforcing our zebrafish *dlc1* mutant as an *in vivo* model to study the etiology of human CHDs.

## RESULTS

### dlc1 and dlc3 are expressed in the lateral plate mesoderm (LPM) and in endothelial cells

Currently, only mouse and chicken models have been established to analyze early *dlc* expression and function during embryonic development^9,10^. To investigate the spatial and temporal distribution of *dlc1, dlc2* (*stard13b*) and *dlc3* during zebrafish development, we performed *in situ* hybridization at different embryonic stages. Specific expression of all three genes was detected from 6 somites (ss) onwards. *dlc1* expression was first detected in the posterior LPM (PLPM) and in the head of the embryo (Fig. 1A, Supplemental Fig. 1A). At 24 hours post fertilization (hpf), *dlc1* expression was broadly distributed throughout embryonic tissues and in the vascular system. This included the mid cerebral vein (MCeV) in the head (Fig. 1B, arrowhead) and the aortic and venous endothelium in the trunk (Fig. 1B, C). *dlc3* was expressed in the anterior LPM and the PLPM (Fig. 1D solid arrows, Supplemental Fig. 1B), intersomitic regions, and the tail bud at 6 ss (Fig. 1D, Supplemental Fig. 1C). At 24 hpf, *dlc3* expression strongly resembled *kdrl* expression, marking the axial trunk vasculature and the first sprouting intersegmental vessels (ISVs) (Fig. 1E). Due to the absence of *dlc2* expression in the LPM and vascular tissues (Supplemental Fig. 1D-G), the physiological role of *dlc2* was not further investigated. The endothelial expression of *dlc1* and *dlc3* was confirmed in sorted *kdr1:eGFP*^***+***^ cells by qPCR analysis. Both genes were enriched in endothelial cells (ECs) at 26 hpf compared to eGFP-negative sorted cells (Fig. 1F, G). Altogether, *dlc1* and *dlc3*, but not *dlc2*, are expressed in putative angioblasts in the LPM and in ECs at later stages of development.

**Fig. 1.**
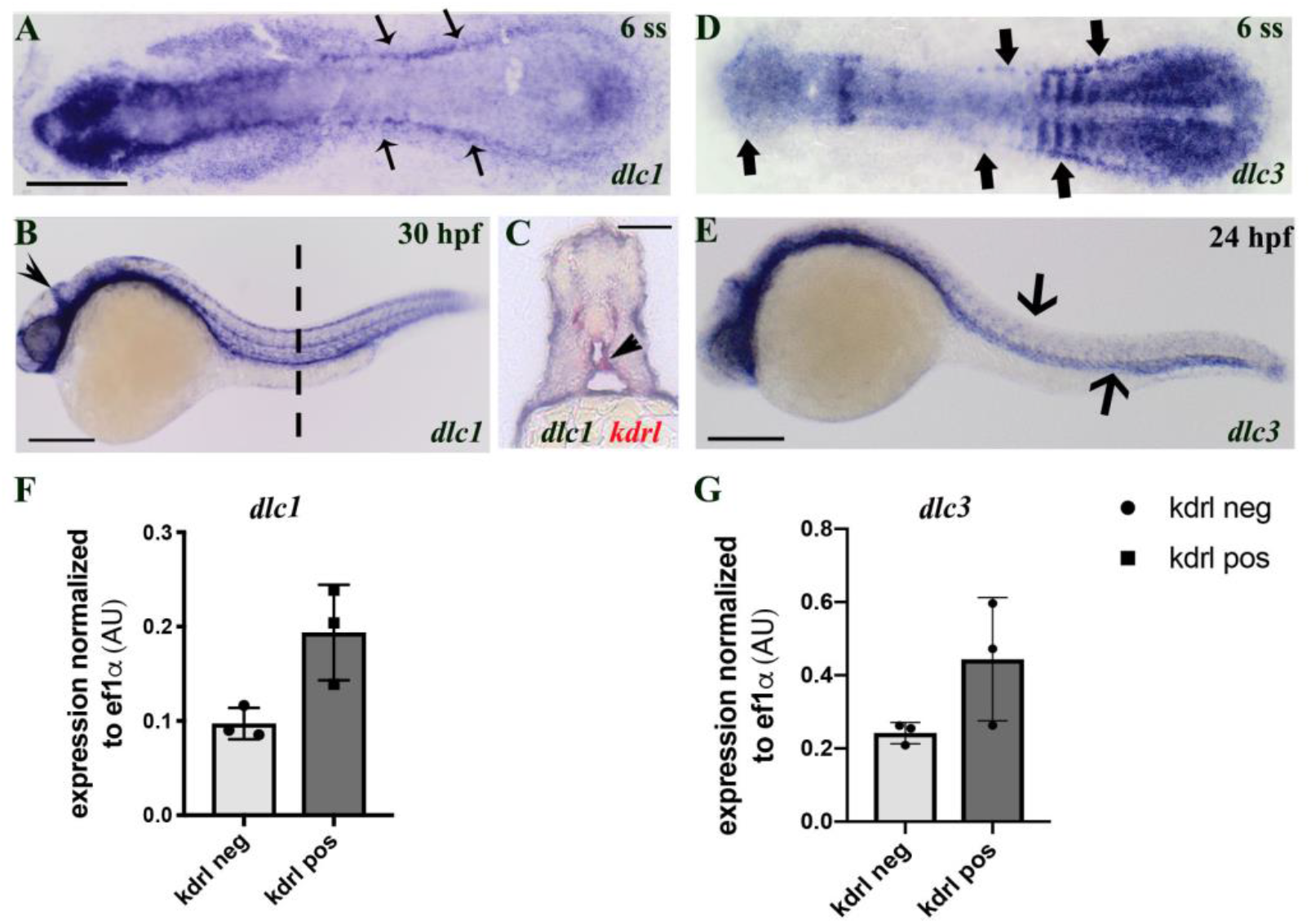
*dlc1* and *dlc3* are expressed in the LPM and in endothelial cells. A) shows a flatmounted ISH image of *dlc1* expression in the PLPM at 6 ss (black slim arrows), Scale bar = 250µm. In B) expression of *dlc1* is shown at 30 hpf (Scale bar = 250 µm). The pointed arrowhead highlights expression in the midcerebral vessels. A corresponding cross-section of the trunk (level of broken line) and double staining of *kdrl* is shown in C) (solid arrowhead), Scale bar = 100 µm. D) and E) show *dlc3* expression at 6 ss and 24 hpf, respectively, Scale bar = 250 µm. F) and G) show endothelial expression levels at 26 hpf (in biological triplicates, N = 3). In all images the embryos are oriented with their head to the left and the tail to the right. All in situ experiments have been performed a minimum of three times (N = 3).

### Overexpression of dlc1 and dlc3 affects angiogenesis

Overexpression of *dlc1* and *dlc3* (by injection of full-length mRNA at the 1-cell stage) induced a high lethality by 8 hpf ranging from 75% to 95% of injected embryos. The surviving embryos showed severe ISV defects, which manifested in hyperbranching and misguided vessels (Fig. 2A-C). To overcome this problem, we established a stable transgenic line, *hsp70:dlc1*, with *dlc1* under the control of the heat-shock promoter *hsp70* (Fig. 2D and Supplemental Fig. 2). Analysis of ISVs at 2 days post fertilization (dpf) following heat shock at 13ss revealed prominent patterning defects. Thus, temporal restriction of *dlc1* overexpression (Supplemental Fig. 2) recapitulated the phenotype observed in mRNA overexpression experiments (Fig. 2D-H), while greatly reducing lethality. Besides the ISV phenotype, heat-shocked embryos also displayed alterations in the growth of the bilateral common cardinal veins (CCVs). The CCV migrates as a collective EC sheet that ultimately connects the venous system and the endocardium of the cardiac inflow tract (IFT). In heat-shocked embryos, the leading front and edges of the migrating CCV cell sheet often appeared uneven and/or small holes were observed in the EC sheet (Fig. 2I, J). These two CCV phenotypes appear to be dependent on the RhoGAP domain of *dlc1*, as truncated *dlc1* mRNAs containing only the GAP and START domains caused a similar phenotype (Fig. 2K, K’), while other truncated constructs of *dlc1* did not induce this phenotype (Supplemental Table 1). When we treated heat-shocked embryos with a Rho Activator, we partially restored the growth of the CCVs (Fig. 2L-O), demonstrating that *dlc1* alters the CCV growth pattern in a GAP-dependent manner.

**Fig. 2.**
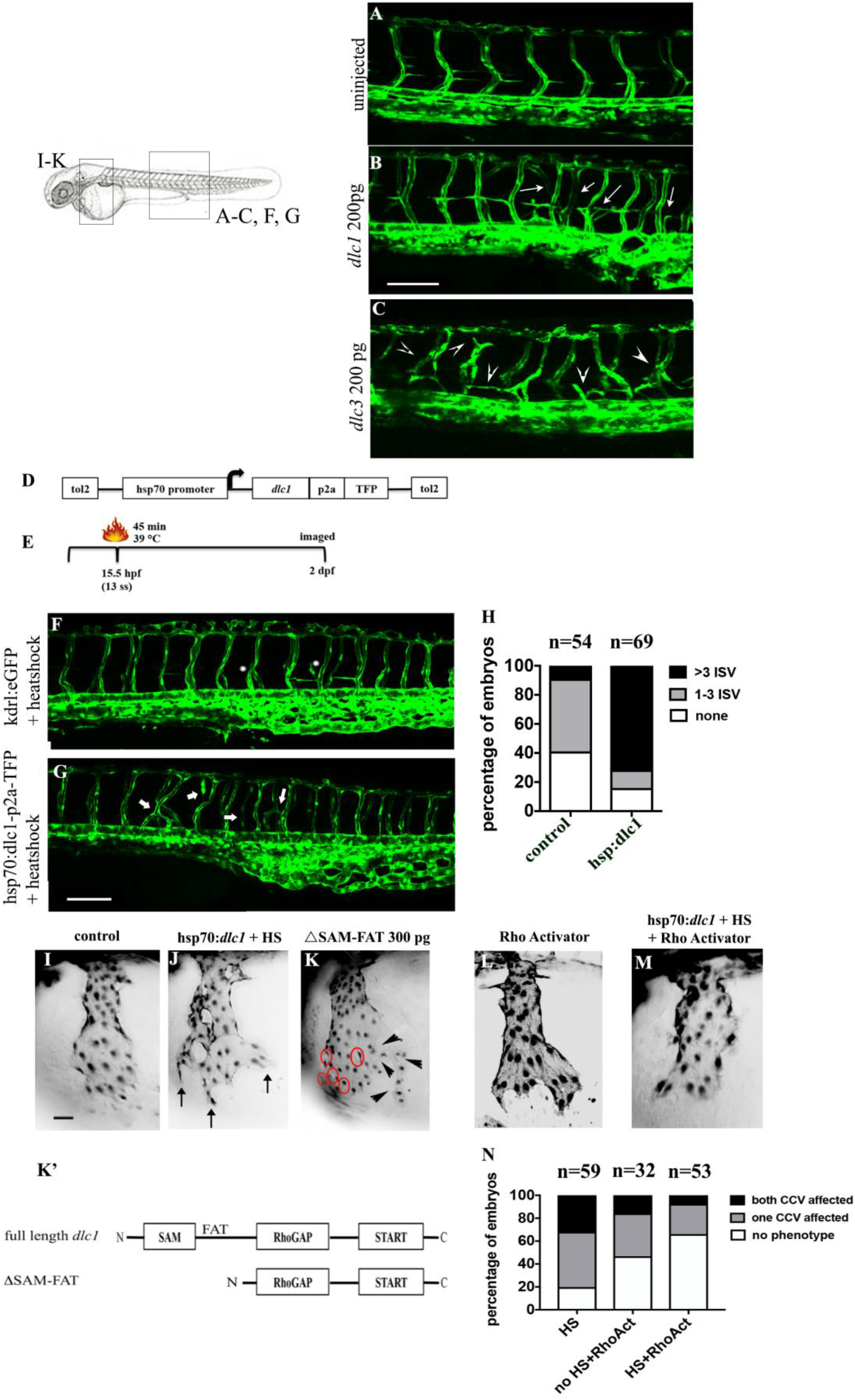
Overexpression of *dlc1* and *dlc3* affects angiogenesis in the trunk and the growth of the common cardinal veins, which is RhoGAP-domain dependent. A) – C) shows an example of *kdrl:eGFP* embryos injected with 200 pg of full length mRNA for *dlc1* (B) and *dlc3* (C), compared to the uninjected control embryo (A), Scale bar = 100 µm. Hypersprouting ISVs are indicated with white arrows (*dlc1*, B) and white arrowheads (*dlc3*, C). D) shows a schematic representation of the *hsp70:dlc1-p2a-TFP* construct and E) the heat shock protocol. Temporal overexpression in an exemplary *hsp70:dlc1/kdrl:eGFP* embryo is shown in G) compared to heat shocked *kdrl:EGFP* control embryos in F) and the corresponding quantification of the ISV phenotype in H), Scale bar = 100 µm. The same heat-shock exemplary phenotypic ISV example image is used in Fig. 2G and Fig. 3N. The ISV phenotype is quantified as either more than 3 ISVs affected (black), 1-3 ISVs (grey) or no ISVs affected (white, none). Counted ISV phenotypes consisted of bifurcations/hypersprouting, additional smaller sprouts and thin/non-lumenized vessels. I) and J) show the CCVs of *hsp70:dlc1/kdrl:eGFP* embryos either non-heat shocked (I) or heat shocked (J), the uneven leading edge is marked by arrows. The occurrence of a phenotype is quantified in the first bar graph in N). The CCV phenotypes were recapitulated after injection of 300 pg of *dlc1* mRNA lacking the SAM and FAT domain (ΔSAM-FAT), only containing the RhoGAP and START domains of *dlc1* (K and K’). The red circles highlight holes in the cell sheet and the solid black arrowheads point to uneven leading edges. L) shows Rho Activator-treated embryos (quantified in the second bar graph in N)) alone. The combined rescue of heat shocked hsp-*dlc1* embryos and Rho Activator treatment is shown in M) (compare with the heat-shock phenotype alone in J)) and is quantified in the third bar graph in N). Scale bar = 50 µm. N) shows the quantification of phenotypes observed in J), L) and M). The quantification factors in the percentages of embryos with both of their CCVs affected (black), only one CCV on one arbitrary side (grey) or no CCV phenotype on either side (white, no phenotype). Any alteration to CCV growth was counted as phenotypic, including holes in the cell sheet and uneven growing fronts/edges. All images show lateral views of the embryo oriented with the head to the left. n indicates the combined total numbers of examined embryos per experimental group from 3 individual experiments (N = 3). The bar graph displays the mean across the 3 pooled experiments.

### dlc1 affects CCV growth via the Sema3d/PlxnD1 and the Nrp1/RhoA axes

The CCV growth pattern induced by *dlc1* overexpression (Fig. 3A) is similar to the phenotype observed in *sema3d* morphants and mutants ^17^. A recent zebrafish study showed that Sema3d regulates collective EC migration in the CCVs by two distinct mechanisms. The guidance and straight outgrowth of the CCV is dependent on Sema3d signaling through PlexinD1, whereas the cytoskeletal integrity of the EC sheet is maintained by Sema3d signaling through Nrp1. The latter regulates actin network organization through RhoA, which stabilizes the EC sheet morphology^17^. To investigate whether *Dlc1* mediates its effect on CCV development via Sema3d and Nrp1, zebrafish *sema3d* and rat *Nrp1* mRNA^17^ were injected in one-cell stage *hsp70:dlc1* embryos, that were then heat-shocked at 13ss. While *sema3d* mRNA rescued the uneven growing edge of the CCV, holes in the migrating EC sheet were still present (Fig. 3B, C, F). Conversely, *Nrp1* overexpression rescued the EC sheet integrity but not the uneven CCV growth pattern (Fig. 3D, E, G). These results suggest that Dlc1 controls EC migration via the Sema3d/PlxnD1 and EC sheet integrity via the Nrp1/RhoA axis during CCV growth.

**Fig. 3.**
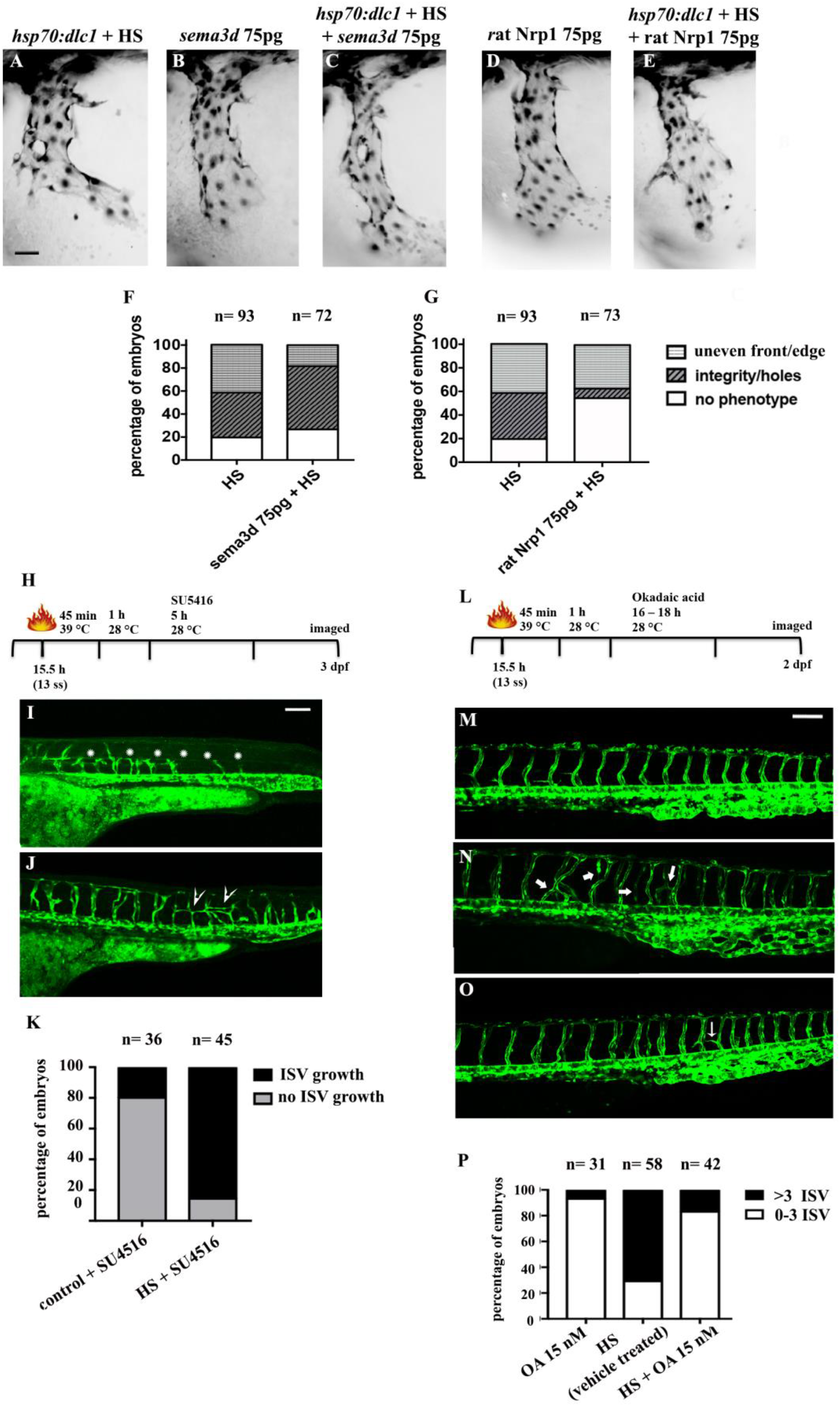
*dlc1* acts downstream of VEGF-A/*kdrl/Nrp1* signaling and its activity is modulated by phosphorylation. Heat shocked hsp70-*dlc1* embryos show a combination of different CCV phenotypes after heat shock (A)-E)), quantified in F) and G)). A) shows hsp70:*dlc1*/*kdrl*:eGFP embryos after heat shock (HS); B) injected hsp70:*dlc1/kdrl*:eGFP with 100pg of *sema3d* mRNA alone (no heat shock) and C) the combination of both, which is quantified in F). D) shows injected hsp70:*dlc1/kdrl*:eGFP with 100pg of rat Nrp1 mRNA alone (no heat shock) and E) the combination of both, which is quantified in G), Scale bar = 50 µm. The graphs in F) and G) distinguish between the two differential CCV phenotypes; displaying the percentages of affected embryos with uneven leading edges in light grey (uneven front/edge; F) or the occurrence of holes in the cell sheet as black diagonal stripes (integrity/holes; G). H) shows the schematic setup of the heat shock experiment followed by subsequent SU5416 treatment. A respresentative image of SU5416 treated embryos on the *hsp70:dlc1/kdrl:eGFP* (control siblings) or *kdrl:eGFP* (wildtype) background that did not receive a heat shock are shown in I) and treated embryos that were additionally heat shocked are shown in J), Scale bar = 100 µm. The percentage of embryos that show recovered ISV growth (black), compared to no ISV growth (grey) is quantified in K). All ISV that did not sprout or failed to sprout beyond the horizontal myoseptum were counted as no ISV growth. L) shows the schematic setup of the heat shock experiment followed by subsequent Okadaic acid (OA) treatment. M) shows a *hsp70-dlc1/kdrl:eGFP* embryo treated with OA, N) *hsp70:dlc1/kdrl:eGFP* embryos that were heat shocked (no treatment) and O) *hsp70:dlc1/kdrl:eGFP* embryos that were heat shocked and received OA treatment., Scale bar = 100 µm. The same exemplary heat-shock phenotypic ISV example image is shown in Fig. 2G and Fig. 3N. The percentage of affected ISV per condition is quantified in P). The ISV phenotype is quantified as either more than 3 ISVs affected (black) or 0-3 ISVs affected (white). Counted ISV phenotypes consisted of bifurcations/hypersprouting, additional smaller sprouts and thin/non-lumenized vessels. Images (I, J, M-O) show lateral views of the embryo with the head towards the left. All experiments have been performed 3 times (N = 3). n indicates the combined total pooled numbers of embryos from the 3 individual experiments above each bar graph. The bar graph displays the means of the 3 pooled experiments.

### dlc1 acts downstream of kdrl signaling

Neuropilins function as co-receptors to enhance the binding activity of Vegfa and Vegfc to Flt1, Kdrl and Flt4, respectively^7,8^. Since we showed that Dlc1 interacts with the Sema3d/PlxnD1 and the Nrp1/RhoA axes, we next examined its role in Kdrl signaling. To investigate this, *hsp70:dlc1* embryos were divided into two groups and only one group was heat-shocked at 13ss prior to treatment with a high dose (1.5µM) of the VEGFR2 inhibitor SU5416 for 5 hours (Fig. 3H). At 3dpf, the non-heat-shocked SU5416-treated embryos showed pronounced vascular defects throughout the whole embryo (Fig. 3I). All vasculogenic and angiogenic sites were affected, which led to a complete absence of blood circulation. In 20% of these embryos, ISVs were able to partially regrow, however, the large majority failed to extend beyond the horizontal myoseptum (Fig. 3I, asterisks). In sharp contrast, temporal-controlled overexpression of *dlc1* restored ISV growth in 83% of examined embryos (Fig. 3J, K). Overexpression of *Dlc1* not only rescued the initial sprouting defect, but also prevented the ISV extension defects that were observed in control SU4516-treated embryos. In line with our initial gain-of-function phenotype, we still observed some additional hyper-sprouting (Fig. 3J, arrowheads). This finding suggests that Dlc1 acts downstream of the Kdrl signaling axis and plays a crucial role during vascular development.

### dlc1 activity is modulated by phosphorylation

In human epithelial cells, DLC1 activity is regulated by a phosphorylation-dephosphorylation cycle downstream of EGFR signaling^18^. Therefore, we investigated whether inhibiting protein phosphatase 2 A (PP2A) dephosphorylation of Dlc1 could suppress its activity *in vivo* (Fig. 3L). Treatment with okadaic acid (OA), a PP2A inhibitor, caused only minimal effects on vascular development (Fig. 3M). *Hsp70:dlc1* embryos were heat-shocked at 13ss and subsequently treated with OA overnight. OA treatment rescued the hyper-sprouting ISV phenotype induced by *dlc1* gain-of-function (Fig. 3N-P), suggesting a conserved regulation of *dlc1* in zebrafish *in vivo*, compared to what was previously observed *in vitro* for human cells.

### The dlc1 mutation impairs the development of the vascular system

By using CRIPSR/Cas9, we generated two different *dlc1* mutant alleles: a -3/+2 bp indel mutation and a -4 bp deletion (Supplemental Fig. 3A). Both mutations resulted in premature stop codons after 174 and 173 translated amino acids, respectively; resulting in severely truncated proteins that lacked most of the FAT domain, and the GAP and START domains. Based on our knowledge of the Dlc family of proteins, we considered these as null alleles. The mutant phenotypes were analyzed on a *kdrl:*eGFP transgenic background from 24hpf until 2.5dpf. Matings of *dlc1*^*+/-*^ adults always induced the same two phenotypes in mutant embryos with a Mendelian ratio (20-25%). However, the distribution of these two phenotypes varied considerably and was independent of the mating pairs. The less severe phenotype consisted of the impaired formation of the lateral dorsal aortae (LDAs) and aortic arches 1 (AA1s) (12%-25% of mutant embryos per clutch), which in most cases did not result in embryonic lethality (Fig. 4). In some cases, the AA1s were functionally impaired, resulting either in stenosis (Fig. 4A, B arrow) or failure to connect to the LDA (Fig. 4B right side, arrowhead). In most cases, the bilateral LDA either failed to correctly merge into one vessel (Fig. 4E, F, F’) or failed to complete proper lumenization (Fig. 4G, G’). This was phenocopied by *dlc1-*MO injection (Supplemental Fig. 3C,D; Fig. 4H, I, I’). Erythrocytes could often only circulate through one LDA, possibly compensating for the stenosis in the other LDA (Fig. 4I arrows). *Dlc1* mutant embryos showed stenotic phenotypes throughout the LDA, including the looped lateral region, where the LDA merge into the AA1 (Fig. 4C, D). Surprisingly, *dlc3* mutants (Supplemental Fig. 3B) showed only transient mild edema and decelerated blood flow, especially in the head, but these phenotypes resolved completely by 2.5dpf (data not shown). Overall, there were no specific changes to the vasculature in *dlc3* mutants that could explain this transient phenotype. Only in rare cases did *dlc3* mutant embryos display a comparable LDA phenotype to *dlc1* mutants (Supplemental Fig. 4A, B). Surprisingly, the *dlc1* or *dlc3* mutants did not show any modified ISV growth, whereas both MOs considerably affected ISV growth in the trunk (Supplemental Fig. 3C-E, Supplemental Fig. 5). *In vivo* time-lapse microscopy of *dlc1* and *dlc3* morphants showed growing ISV stalling at the horizontal myoseptum, as they failed to properly elongate dorsally (Supplemental Fig. 5A-F and Supplemental Movies 1-3). To determine if the *dlc1* morphants showed an alteration in tip and/or stalk cell identity, we purified ECs from their trunks and tails (Supplemental Fig. 5G). There were no changes in the expression of genes involved in tip or stalk cell identity between *dlc1* morphants and control embryos (Supplemental Fig. 5H). This indicated that *dlc1* likely controls the directed EC migration of sprouting ISVs but does not affect the identity of tip and stalk cells in the angiogenic sprout. Taken together this data suggests that *dlc1* controls vasculogenesis in the LDA and AA1s and EC migration during angiogenic sprouting in zebrafish.

**Fig. 4.**
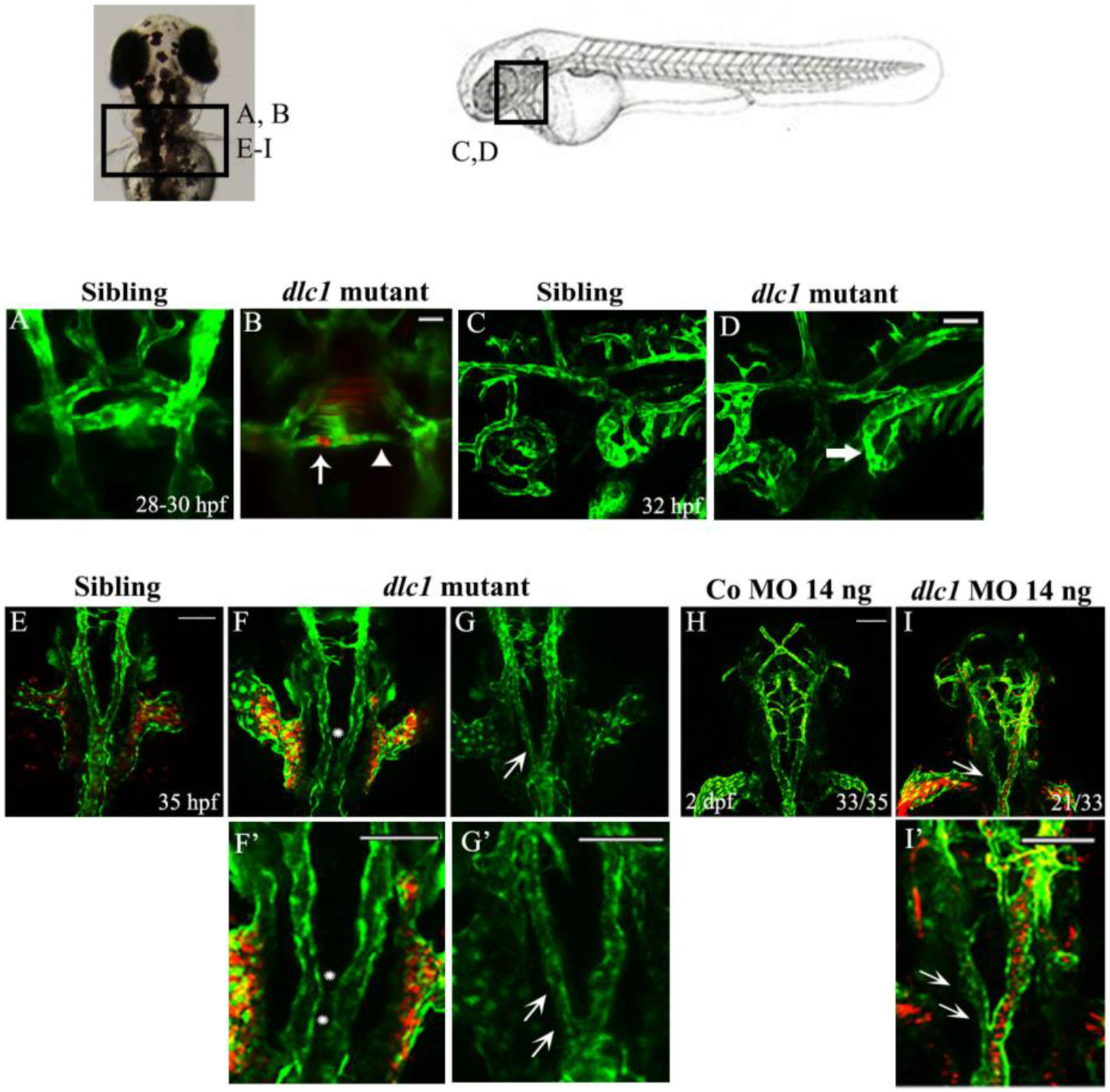
Loss of *dlc1* affects the formation of the LDA and AA1s. A) shows the AA1 of WT/siblings on a *kdrl:eGFP* background (dorsal view). B) shows the *dlc1* mutant with red blood cells (*gata1:dsRed*) trapped in the AA1 (arrow, left) and a stenotic/partially unconnected AA1’ (arrowhead, right) at 28-30 hpf, Scale bar = 25 µm. C) and D) show lateral views of the looped connection of the LDA to the AA1 in control siblings (C)) and in *dlc1* mutants (D), stenosis indicated with a solid arrow) at 32 hpf, Scale bar = 50 µm. E) to I) show dorsal views of the LDA. F) and G) show LDA phenotypes in *dlc1* mutants, compared to WT siblings in E) at 35 hpf. The differential mutant LDA phenotypes consist of not properly fused LDA into the single dorsal aorta (asterisk, F) or stenotic LDA (arrow, G) without blood flow. The panels F’) and G’) show magnifications of the observed phenotypes. All mutant phenotypes have been observed across several individual experiments (n ≥ 5) in 1 - 4 embryos per experiment. I) shows a similar LDA phenotype after *dlc1* MO injection, compared to control MO injected embryos in H) at 48 hpf, Scale bar = 100 µm. The numbers in H) and I) demonstrate the frequency of embryos with the indicated phenotype from one exemplary injection experiment. The panel I’) shows a magnification of the observed phenotype. MO injection experiments have been performed at least 3 times. The images B), E), F) and I) include the *gata1:dsRed* transgene to highlight the presence or absence of blood flow.

### dlc1 mutant embryos as a model of congenital heart defects (CHDs)

A second phenotype was found in 0%-13% of embryos obtained from *dlc1*^+/-^ incrosses, which did not survive past 5dpf (Fig. 5). This severe phenotype was observed as early as 28-30 hpf due to the lack of blood circulation and the development of pericardial edemas (Fig. 5A, B). Microangiographies confirmed that no blood or liquid could leave the heart and enter circulation in these mutants (Fig. 5C, D). The absence of blood circulation was mainly due to a defective immature outflow tract (OFT; 80%), resulting in backflow of blood from the ventricle to the atrium (Fig. 5B, Supplemental Movies 4 and 5), preventing blood from leaving the heart. Furthermore, the defective OFT formation resulted in enlargement and dilation of the heart but did not affect atrium and ventricle specification (Fig. 5E, F). qPCR analysis on pools of isolated mutant hearts revealed significantly higher expression of *plxnd1* and a trend towards elevated *kdrl* expression (not significant) compared to control hearts, whereas expression of the cardiomyocyte marker *myl7* was similar (Fig. 5G). Due to the cardiac enlargement, the observed transcriptional changes could either be attributed to more endocardial cells or to an overall higher expression of *plxnd1* and *kdrl*. Moreover, mutant hearts displayed a change in *bmp4* expression that normally marks the OFT, the atrio-ventricular (AV) canal, and to a much lesser extent, the IFT (Fig. 5H, M). In *dlc1* mutants, *bmp4* staining was highly present in the OFT and the IFT, but only remotely in the AV canal (Fig. 5I, M), even though no changes in overall *bmp4* expression could be detected by qPCR (Fig. 5J). This rather suggests a change in *bmp4* distribution within the heart than a change at the transcriptional level (Fig. 5M). *dlc1* morphants phenocopied the abnormal *bmp4* distribution in the heart (Fig. 5K, L), but not the obstructed OFT phenotype. This indicates that the changes in *bmp4* distribution are probably not a result of the morphological alterations of the heart due to the lack of blood flow, but rather depended on the proper migration of *bmp4*-expressing cells within the heart of *dlc1* mutants. Decreased *dlc3* levels did not cause any cardiac phenotypes, but *dlc1*/*dlc3* double mutants exhibited higher occurrence of the cardiac phenotype. Concordant with this hypothesis, double mutant fish never survived to adulthood. Moreover, injection of low doses of the *dlc3*-MO (4 ng) into *dlc1* mutants favored the appearance of the severe cardiac phenotype by 2-fold (Fig. 5N). These results point towards a certain degree of redundancy between *dlc1* and *dlc3* during cardiovascular development, although *dlc1* seems to be the major player in this process. Overall this suggests that *dlc1* is indispensable for initial early OFT formation and proper cardiogenesis in the zebrafish embryo.

**Fig. 5.**
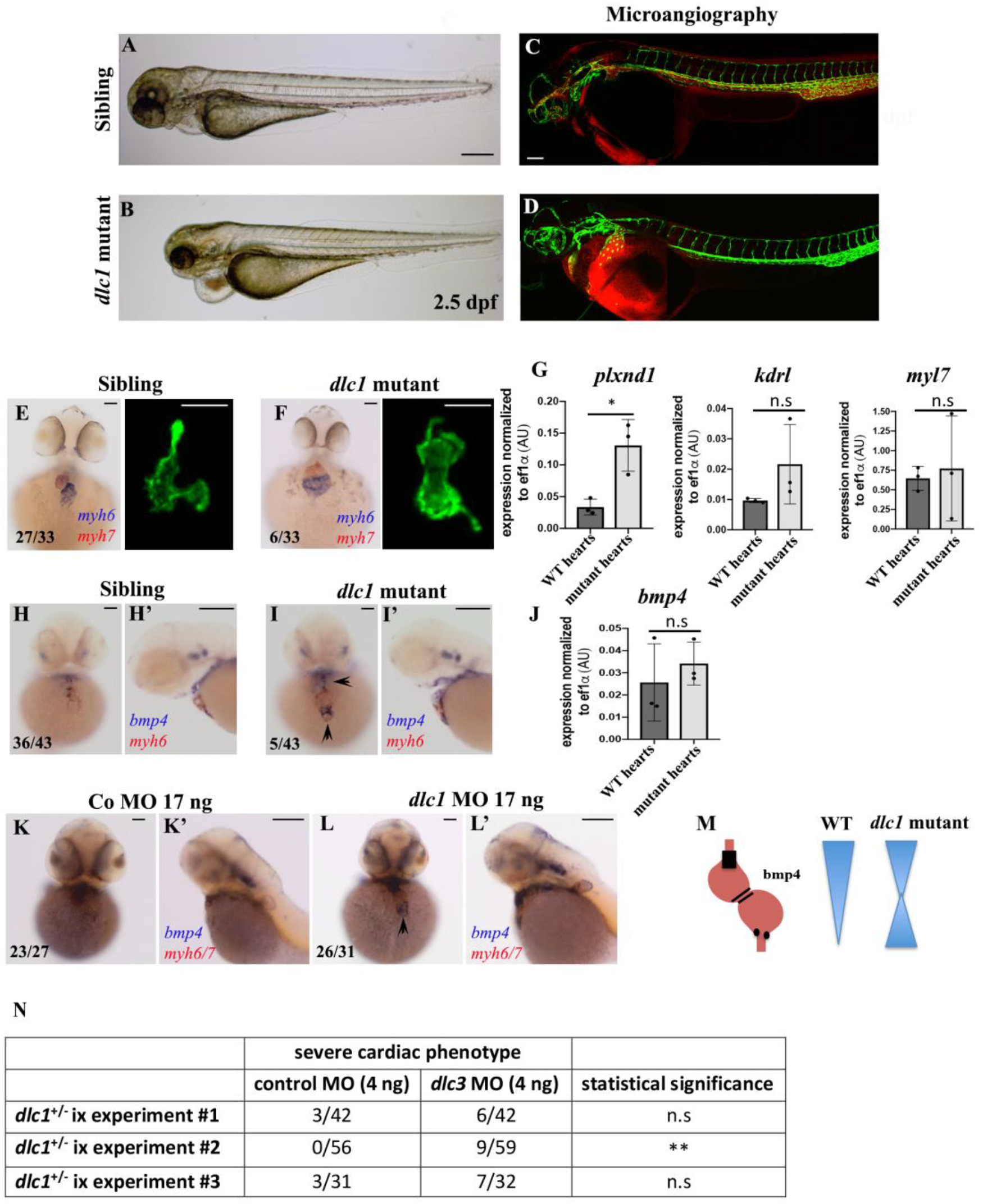
The severe phenotype in the *dlc1* mutant as a model of congenital heart defects. A) – B) show brightfield images of control siblings (A)) and *dlc1* mutants (B)) with the severe heart phenotype, Scale bar = 250 µm. C) and D) shows microangiographies with Rhodamine-Dextran (red) for C) control embryos and D) *dlc1* mutants on a *kdrl:eGFP* transgenic background, Scale bar = 100 µm. C) and D) show exemplary images from one experiment, overall the experiment has been performed on n = 4 different mutant embryos in individual experiments (N = 3). E) and F) show ventral views of ISH for the heart ventricle (blue, *myh6*) and the atrium (red, *myh7*). The numbers in the ISH image display the frequency of embryos with the indicated staining from one exemplary heterozygous in-cross experiment. Next to the ISH images, isolated hearts on a *kdrl:eGFP* transgenic background of control siblings (E) and *dlc1* mutants are shown with altered heart morphology (F). Scale bars = 100 µm. G) shows qPCR analysis of endocardial (*plxnd1, kdrl*) markers and a cardiomyocyte (*myl7*) marker in WT hearts compared to mutant hearts (grey). H) and I) show ventral and H’) and I’) lateral views of *myh6* (red) and *bmp4* (blue) expression in control siblings (H, H’) and *dlc1* mutants (I, I’). H) and I) Scale Bar = 100 µm, H’) and I’) Scale Bar = 500 µm. qPCR analysis for *bmp4* expression in isolated hearts is shown in J). Statistical significance for qPCR (G and J) was determined using an unpaired, 2-tailed *t* test. *plexind1* p = 0.017 (*p < 0.05, significant); *kdrl* p = 0.189 (n.s); *myl7*: p = 0.767 (n.s); *bmp4*: p = 0.501 (n.s). All bar graphs show the mean ± SD of 3 independent experiments (N = 3). The mean consists of 5 -10 hearts per individual biological group (n = 22 pooled hearts in total). K) and L) show ventral and K’) and L’) lateral views of *myh6/7* expression (red) and *bmp4* (blue) in control MO (K, K’) and *dlc1* MO (L, L’) injected embryos. The arrowhead highlights increased staining/expression. K) and L) Scale bar = 100 µm, K’) and L’) Scale bar = 500 µm. M) shows a schematic cartoon of the changed *bmp4* distribution in *dlc1* mutants hearts compared to WT expression. N) shows the increased numbers of *dlc1* mutants with the severe phenotype after *dlc3* MO injection compared to control MO injected embryos, statistical significance for N) was analyzed using Fisher’s exact test. The second experiment proved to be statistically significant (**p < 0.01) with p = 0.0029. For N) one clutch of embryos from a single adult mutant pair was divided equally for control MO and *dlc3*-MO injection, respectively.

## DISCUSSION

Our work represents the first detailed *in vivo* characterization of *dlc1* and *dlc3* in a vertebrate model that can survive several days despite severe cardiovascular phenotypes. While many excellent cell culture studies provided topical and cancer-related pathophysiological data on Dlc1’s functions, they ultimately lack the anatomical context of a whole animal. This makes it difficult to fully understand the physiological role of the Dlc1 protein, in particular when neighboring tissues and cell complexes are missing to provide a contextual scaffold. This in turn may influence the occurrence and consequential manifestation of multifaceted and/or spatially distinct phenotypes.

We showed that *dlc1* and *dlc3* are initially expressed in the LPM and become enriched in ECs during zebrafish development. The overexpression of DLC1 in human cells results in enhanced velocity of migrating cells and a loss of their directionality due to dephosphorylation of focal adhesion proteins^19^. We found that those alterations of cell migration translated into differential phenotypes at two distinct anatomical sites in the zebrafish. While gain-of-function greatly affected angiogenic sprouting in the zebrafish trunk, causing mis- and hyperbranching ISV growth, we also observed an uneven leading edge of ECs, as well as a disrupted EC layer during CCV growth. The specificity of the trunk phenotype was confirmed by rescuing the aberrant ISV sprouting phenotype through inhibiting the final activation step of Dlc1 using a specific PP2A inhibitor. This mechanism was previously described by Ravi and colleagues^18^ in epithelial cell culture studies and appears to be globally conserved *in vivo*.

In zebrafish, the loss of directional vessel growth is associated with deficient Sema3a/PlxnD1 signaling during trunk angiogenesis^20^ and this pathway directly interacts with the Vegfa/Kdrl/Nrp1 axis^21,22^. We found that *dlc1* overexpression can prevent the arrest of ISV growth induced by Vegfr2 pathway inhibition, suggesting that Dlc1 acts directly downstream of the Kdrl axis *in vivo*. In addition to ISV hyperbranching following *dlc1* overexpression, we observed altered migration patterns of ECs during CCV growth. This phenotype has been described in *sema3d*-deficient zebrafish embryos^17^, as Sema3d regulates CCV growth in a *plxnd1*-dependent manner^17^. Furthermore, Sema3d undergoes EC-specific autocrine signaling through Nrp1 to modulate actin network organization via RhoA and ROCK^17^, which is important to preserve the EC sheet integrity. In line with this study, we showed that *sema3d* mRNA injection exclusively rescued the uneven growth of the front and edges of the CCV, while rat *Nrp1* mRNA solely rescued the integrity of the endothelial layer in temporally controlled *dlc1* overexpression experiments. During angiogenesis, attractive and repulsive signaling cues need to be translated into cytoskeletal rearrangements. NRP1-knockdown *in vitro* disrupts F-actin network organization in HUVECs^23^, suggesting a direct interaction of NRP1 and RHOA-mediated actin cytoskeletal rearrangements. Consequently, overexpression of truncated *dlc1* mRNA encoding the GAP and START domains was sufficient to induce a similar, but milder CCV phenotype, compared to full-length *dlc1* overexpression. This indicates that the disruption of the straight growing edge and lesions in the EC sheet are a result of Dlc1 GAP-dependent inactivation of RhoA, which could be partially rescued by a RhoA activator. These results support the role of Dlc1 in orchestrating EC migration via the Sema3d/Plxnd1, as well as the Nrp1/Rhoa axis during CCV growth in zebrafish. We propose that *dlc* RhoGAPs may represent a functional link in translating pro-(VEGF) and anti-angiogenic (Sema3/Nrp1) guidance cues into cytoskeletal rearrangements *in vivo*. However, it is still unclear how *dlc1* precisely affects single components of the actin cytoskeletal machinery *in vivo* during angiogenesis, and how it interacts with the Sema3/Nrp1 axis.

Our results confirm that Dlc1 activation requires dephosphorylation *in vivo*, which is consistent with *in vitro* results that showed ERK-mediated phosphorylation-dephosphorylation cycles are essential for DLC1 to become fully functional^18^. ERK inactivation of FAK triggers the release of the activated FAK binding complex. This enables PP2A-mediated dephosphorylation of the Ser-(P)308 and Thr-(P)301 residues in DLC1, leading to its activation^18^. FAK represents an important component in this signaling cascade and it can also be inactivated following dephosphorylation by Sema3E *in vitro*, thereby modulating anti-angiogenic signaling^24^. We propose that Dlc1 acts downstream of the Vegfa/Kdrl/Nrp1 signaling cascade (Fig. 6), while undergoing the phosphorylation-dephosphorylation cycle and subsequently inactivating RhoA, which was previously described *in vitro*^18^ *(Fig. 6). Moreover, our data also suggests that dlc1* acts downstream of the Sema3/PlxnD1 axis (Fig. 6), thereby providing a functional link between guidance cues and cytoskeletal rearrangements to control directed EC migration *in vivo*.

**Fig. 6.**
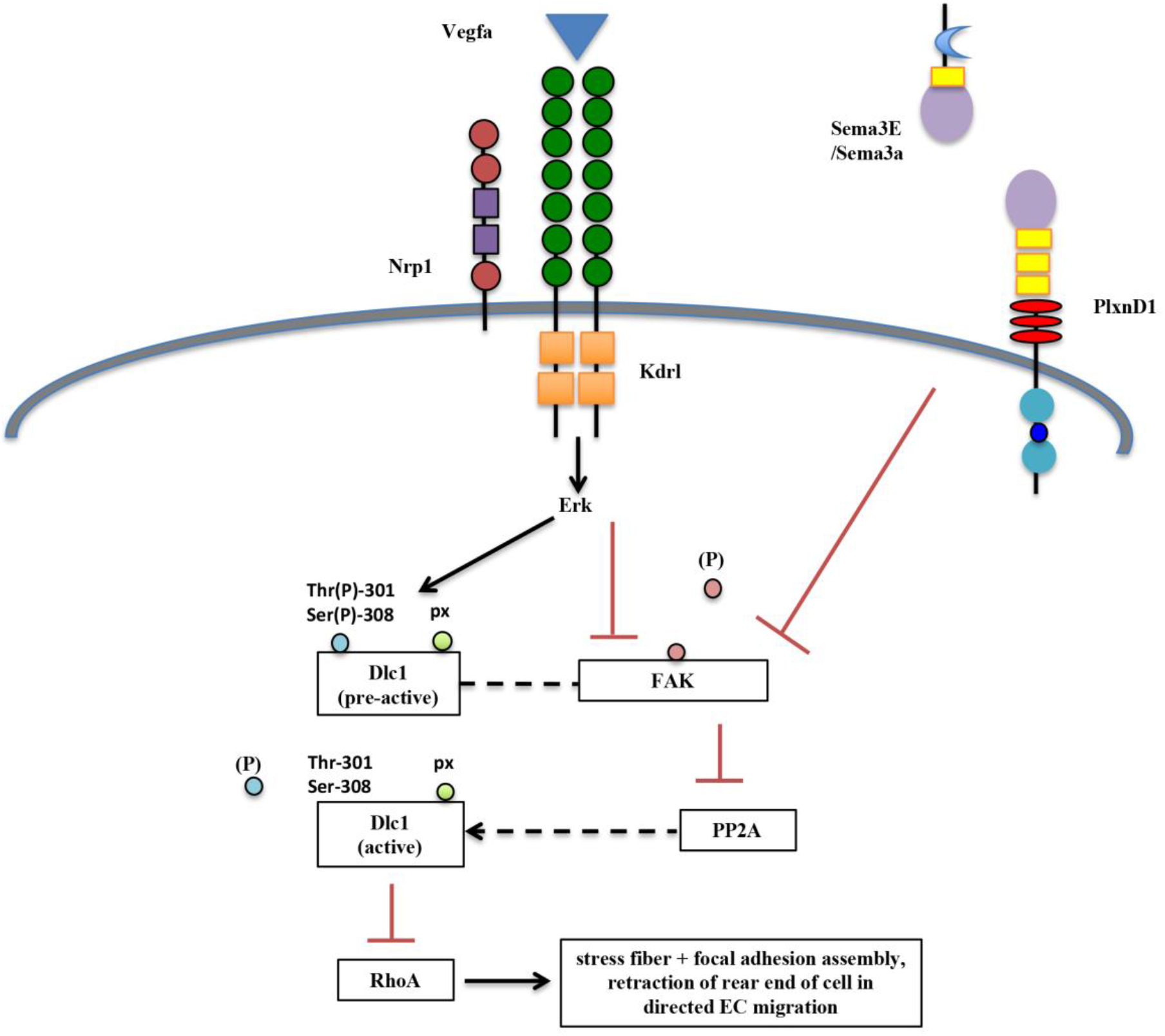
Proposed signaling pathway for *dlc1* in ECs during *in vivo* vascular development. Activation of the VEGF-A/*kdrl/Nrp* axis results in ERK activation that is important for inducing a pre-active state of *dlc1*. After inactivation of FAK and subsequent release of the bound pre-activated *dlc1*, PP2A can dephosphorylate *dlc1* on specific residues and fully activate *dlc1*, which in turn deactivates RhoA. Inactivation of RhoA causes stress fiber and focal adhesion disassembly, which controls the retraction of the rear end of endothelial cells during cell migration. *Sema3E/PlxnD1* signaling has been shown to inactivate FAK, providing a link of the VEGF-A/*kdrl/Nrp1/dlc1* axis and the *Sema3E/PlxnD1*/FAK axis.

Proper ISV growth was severely impaired in *dlc1* and *dlc3* morphants and diametrically opposite to the gain-of-function phenotype. Time-lapse analysis showed that *dlc1*-deficient embryos displayed abnormal ISV growth including stalled sprouts, extension defects, and in severe cases, the formation of vascular plexi instead of elongated vessels. As tip and stalk cell identities appeared normal in *dlc1* morphants, an explanation for this phenotype could be the aberrant function of repellent guidance cues. A stalling phenotype for growing ISVs has also been described in global *sema3d*-KO experiments^17^. However, further investigation is needed to identify which Class 3 Semaphorin is involved in conjunction with PlxnD1/Nrp1 in *dlc* morphants. While ISV development was impaired in *dlc1* morphants but not in mutants, we believe several factors could potentially be responsible for this variance. One explanation could be a partial genetic compensation by other *dlc* family members or even other genes. Another reason could be the fact that proper angiogenesis and lumenization of vessels depend on sufficient blood flow and shear forces, both of which are impaired in several circumstances of the observed *dlc1* mutant phenotypes. Those alterations might mask the emergence and penetrance of an ISV phenotype in *dlc1*^*null*^ embryos.

Besides its role in angiogenesis, knockout of *dlc1* impaired proper AA1/LDA patterning as well as OFT development. AA1/LDA stenosis could be linked to a defective epithelial-to-mesenchymal transition (EMT), a process that converts immobilized ECs into highly motile cells that are able to move through the extracellular matrix. EMT is crucial in the development of most adult tissues and organs^25^, especially the heart^26^. It involves tight regulation of differential expression and/or activation of RhoGTPases^27^, among others. Effects on proper EMT has been associated with ectopic DLC1 expression in nasopharyngeal carcinoma cells ^28^, it is plausible that a similar mechanism could contribute to proper AA1/LDA patterning and OFT development in the zebrafish. Defects in OFT development are the cause in 30% of congenital heart defects (CHD) in humans^29^. Similarly, we showed that the formation of the immature cardiac OFT is defective in *dlc1* and *dlc1/dlc3* double zebrafish mutants resulting in embryonic lethality around 5dpf. Cardiac morphogenesis has been shown to be dependent on RhoGTPase signaling and proper EMT in chick and mice^30,31^. Furthermore, mutations in PLXND1 are associated with truncus arteriosus in humans, which was recapitulated in *PlxnD1*^*null*^ mice that die shortly after birth^22,32^. These mice exhibit defective aortic arch arteries, persistent truncus arteriosus and septal defects^33^. Additionally, this cardiac phenotype is dependent on the GAP domain of PlxnD1 and similar defects have been observed in *Sema3C* and *Nrp1*^*null*^ mice^34-35,36^. Plexins have previously been shown to bind RhoGTPases^37^, it is possible that RhoA and its regulators, such as Dlc1, could be present nearby to transduce signals to the actin cytoskeleton to direct EC migration and EMT.

Isolated hearts from *dlc1*^*null*^ zebrafish mutants showed increased *plxnd1* and *kdrl* expression compared to WT hearts, whereas *myl7* remained largely unchanged. The overall expression level of *bmp4* was unaltered in mutant compared to WT hearts, however there was a striking change in its expression pattern. The alterations in the *bmp4* expression pattern in *dlc1*^*null*^ embryos could point towards an impaired migration of *bmp4-*expressing cells in the *dlc1*^*null*^ zebrafish heart. It is unlikely that *bmp4* distribution was affected by the abnormal heart shape as *dlc1* morphants displayed the same altered *bmp4* expression pattern, without showing the OFT defects seen in mutants. It is very likely that the observed AA1 phenotypes and the disrupted OFT development are interdependent, as a recent study showed that ECs from the aortic arches, in particular the AA1, migrate and contribute to OFT formation^38^. However, attempts to image OFT and AA1/LDA formation using time-lapse microscopy were prevented by the delayed folding speed of the eGFP fluorophore in our transgenic lines and the depth of the immature OFT within the tissue at 20-22 hpf. Future studies of the earliest events in OFT formation would benefit from the generation of transgenic *dlc* mutants with fast superfolding enhanced fluorophores^39^ and the use of specialized microscopy equipment, such as the light sheet fluorescence microscope or SPIM.

The role of *dlc3* during cardiovascular development remains elusive as no cardiac phenotypes were observed in mutants. However, our data suggest a partial redundancy with *dlc1*, while *dlc2* seemed not involved in this process. Future experiments should address potential compensatory mechanisms between the three family members. It is noteworthy that the zebrafish *dlc1* mutant displays a causative phenotype of CHD, which is congruent with the enrichment of rare DLC1 variants in human CHD patients^15^. As genetic models of sporadic CHD are rare and the *Dlc1* mutation causes early intrauterine embryonic lethality in the mouse model^9^, the zebrafish *dlc1* mutant will provide a valuable tool to study the underlying mechanisms involved in early OFT development.

## MATERIALS AND METHODS

### Zebrafish husbandry and maintenance

Zebrafish procedures were approved by the Geneva Veterinary Office and were performed according to the guidelines from Directive 2010/63/EU of the European Parliament. All experiments were performed using the AB^*^ wildtype strain and the following transgenic animals: Tg(*kdr1*:eGFP)^s843 40^, Tg(*gata1*:dsRed)^sd2 41^, Tg(*fli1a:lifeact*-GFP)^mu240 17^ and Tg(*myl7*:dsRed)^s879 42^. Embryos were dechorionated at 24 hpf and incubated with 0.002% PTU to prevent pigment formation. Embryos and larvae were anaesthetized with 0.2 mg per mL of Tricaine methanesulfonate (MS-222) in E3 medium by immersion. For fixation, the embryos and larvae (older than 24 hpf) were first anaesthetized with Tricaine and subsequently fixed in 4% cold paraformaldehyde (PFA).

### mRNA and Morpholino injections

mRNA was transcribed using the mMessage mMachine Kit SP6 (Ambion) according to the manufacturer protocol from a linearized pCS2 vector containing the open reading frame of the polymerase chain reaction amplified product. After transcription, the mRNA was purified by phenol-chloroform extraction and resuspended in RNAse-free water. 1 nL containing 100-200 pg of full length mRNA or the stated concentration of the respective Morpholinos (MO, purchased from Gene Tools) were injected into 1-4 cell stage embryos. *dlc1*-MO 5’-AGTCTTCACTCTGAAACGATATGGA-3’, *dlc3*-MO 5’-CTCTGCTGATGGTAACAGACACAGA-3’, standard control MO, 5′-CCTCTTACCTCAGTTACAATTTATA-3′.

### Generation of transgenic lines (Tol2) and mutant lines (CRISPR/Cas9)

For Tg(*hsp70l:dlc1-p2a-TFP*) fish generation, the sequence for the zebrafish *dlc1* (without the STOP codon), was inserted in a Tol2 vector containing the *hsp70l:xxxp2a-TFP* sequence (kind gift from Prof. B. Martin, Stony Brooks University, NY) by In Fusion (Clontech) cloning. For transgenesis, 1-cell stage embryos were injected with 30 pg of the final Tol2 vector, along with 30 pg Tol2 mRNA.

For CRISPR/Cas9-mediated generation of *dlc1* and *dlc3* mutants, single-guide RNA was generated by annealing oligonucleotides at 95°C for 5 min, and then 22°C for 45 min. The samples were diluted 20-fold and ligated into the pDR274 plasmid (kind gift from Keith Joung; Addgene plasmid 42250), which was previously linearized with BsaI. Guide RNA was generated using the MEGAshortscript T7 kit (Ambion). 500 pg recombinant Cas9 protein (PNAbio) and 250 pg of the guide RNA were co-injected per embryo. The guide RNA sequences for *dlc1* were: 5’-GGACCTTAGCGAGCAGCCAG-3’ and for *dlc3*: 5-CAGGCAACCGCTCCTCAACC-3’. Embryos were screened using primers that flanked the target region (Supplemental Table 2), and mutant alleles identified by sequencing.

### Whole-mount in situ hybridization

Whole-mount *in situ* hybridization (WISH) was performed on 4% PFA-fixed embryos as previously described^43^. In brief, RNA probes were generated by linearization of TOPO-TA vectors (Invitrogen) containing the PCR-amplified cDNA sequence. Digoxigenin- or FITC-labeled antisense probes were synthesized using an RNA Labeling Kit (SP6/T7; Roche). The staining was revealed with NBT/BCIP or INT/BCIP substrate (Roche). Embryos were imaged in 100% glycerol, using an Olympus MVX10 microscope. Oligonucleotide primers used to amplify and clone cDNA for the production of WISH probes are listed in Supplemental Table 2.

### Cryosections

Fixed embryos were subjected to WISH staining and washed afterwards in methanol, followed by PBS. Embryos were mounted in OCT and frozen at -20°C. 5 µm thick sections were obtained using a Cryostat, the consecutive sections were then transferred to Superfrost Microscope slides and stored at -20°C. Sections were defrosted and mounted in Glycerol shortly before imaging on a Zeiss Axioskop 2 plus.

### Heat shock and chemical inhibitor treatments

Heat shock was performed at the 13-somite stage (ss) of embryonic development for 45 min in a 39°C water bath. Embryos were returned to 1xE3 (room temperature) and incubated at 28°C. Subsequent chemical inhibitor treatments were performed 0.5-1 hour after the heat shock and incubated for the stated times at 28°C in the dark. The chemicals were diluted in 1xE3 in the following concentrations: 1.5µM of the *VEGFR2* inhibitor SU5416 (Selleckchem) and 15nM Okadaic acid (Enzo).

### Confocal microscopy

Live (anaesthetized) embryos were embedded in 0.5% agarose supplied with 0.07 mg per mL Tricaine in E3 in a glass-bottom dish. Confocal imaging was performed using a Nikon inverted A1r spectral. For time-lapse experiments, embryos were imaged in a heated chamber at 28°C overnight. The larvae were kept anaesthetized for the duration of the experiment (several hours to overnight). Images were analyzed and the contrast and brightness equally adjusted using the FIJI (ImageJ) software.

### Microangiography

3-4 nl of 2 mg per mL Rhodamine B isothiocyanate-Dextran (70kDa, Sigma) dissolved in PBS and 5 mM HEPES were injected into the sinus venosus of 30 hpf embryos. For this, the embryos were anesthetized with Tricaine, placed dorsally into an injection mold and fixed in place by applying 3% Methylcellulose. Embryos were immediately imaged after the injection (≤ 30 min).

### FACS

Embryos were dissociated in 0.9x ice-cold PBS using 0.5mg per mL Liberase (Roche) then resuspended in 0.9x PBS/1% FCS. Dead cells were excluded by SYTOX-red (Life Technologies) staining. Cell sorting was performed using an Aria II (BD Biosciences). Sorted cells were collected in RLT buffer (Qiagen) and frozen at -80°C.

### Heart isolation

Isolation of embryonic hearts was performed in general as previously described^44^ with slight adjustments. Batches of not more than 20 embryos were anaesthetized in Tricaine/E3 and washed three times with ice cold Leibovitz’ L15 medium (Invitrogen) supplemented with 10% FCS. The embryos were then mechanically dissociated (20-30x) in L15 medium/10% FCS using a syringe and 19G needle mounted to a ring stand and directly transferred to a petri dish containing the same medium. Intact isolated hearts were manually collected with a P20 pipette (5-10 hearts per pool) and immediately placed on ice. Identification of the hearts was achieved by a combination of DIC microscopy and fluorescently marked hearts in the transgenic lines *kdrl*:eGFP (endocardium) or *myl7*:dsRed (cardiomyocytes). After mechanical dissociation of the hearts in 0.9x PBS using a 27G needle, heart cells were recovered and resuspended in RLT buffer.

### RNA extraction, cDNA synthesis and quantitative real-time PCR

RNA was extracted using the RNeasy Minikit (Qiagen) according to the manufacturer protocol and reverse transcribed into cDNA using the Superscript III kit (Invitrogen) or the qScript Kit (Quantabio). Quantitative real-time PCR (qPCR) was performed using the KAPA SYBR FAST Universal qPCR Kit (KAPA BIOSYSTEMS) in a QuantStudio3 system (Life Technology). All primers used are listed in Supplemental Table 2.

### Statistical analysis and Reproducibility

Each experiment has been performed at least 3 times on different experimental days (biological replicates; N). Embryos were individually assessed on the day of the experiment and the data pooled for graphical and statistical analysis. The total and pooled numbers of embryos within a full set of experiments (N = 3) was defined as n. Specific n values are presented above bar graphs per condition where applicable. All data in graph bars represent the mean ± SD. Stacked bar graphs containing multiple groups display only the mean for clarity. Statistical significance of differences between 2 groups was determined using an unpaired, 2-tailed *t* test, while the significance of multiple groups was determined using a 1-way ANOVA with Dunnett’s post hoc correction. To analyze the independence of the occurrence of the *dlc1* mutant phenotype after *dlc3* MO injection, the Fisher’s exact test was used. P values of less than 0.05 were considered statistically significant. All statistical analyses were performed using GraphPad Prism 8.0 (GraphPad Software).

## Data availability

The datasets generated during and/or analysed during the current study are available from the corresponding author on reasonable request.

## Acknowledgements

JYB was founded by the Swiss National Fund (#31003A_166515), and was endorsed by a Chair in Life Sciences from the Giorgi-Cavaglieri Foundation. We would like to thank Chantal Combepine and Corentin Pasche for excellent technical assistance, as well as Dr. W. Herzog and Dr. Lucia Du for critical reading of the manuscript.

## Author contributions

TL executed the experimental work. JYB designed the experiments, provided funding and supervised the research. TL and JYB jointly wrote the manuscript.

