## supplementary data for "*Dlc1* controls cardio-vascular development downstream of Vegfa/Kdrl/Nrp1 signaling in the zebrafish embryo"

### SUPPLEMENTAL FIGURES

#### Supplemental Fig. 1

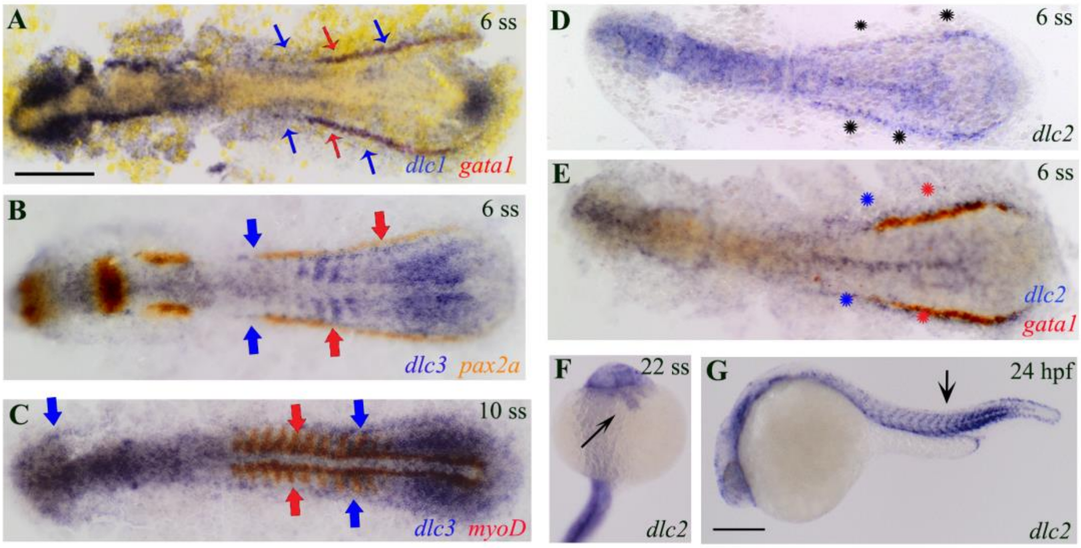

**Suppl. Fig. 1 *dlc1* and *dlc3* are expressed in the LPM and in endothelial cells, whereas *dlc2* is expressed in the intermediate mesoderm**

A) and B) show flatmounted double ISH images of *dlc1* expression in the PLPM at 6 ss (blue slim arrows) and *gata1* expression in red (red slim arrows). In B), expression of *dlc3* is shown in the PLPM at 6 ss in blue (blue solid arrows) and *pax2a* staining in the intermediate mesoderm in red (red solid arrows). C) shows double staining of *dlc3* in blue and *myoD* in red. D) shows *dlc2* expression in blue at 6 ss and double staining with *gata1* in red in E) (asterisks in black (D) and with respective colors in E)). At 22 ss, *dlc2* shows weak expression in the heart tube (slim black arrow, F)) and somitic staining at 24 hpf in G). In all images the embryos are oriented with their head to the left and the tail to the right.

**Supplemental Fig. 2**

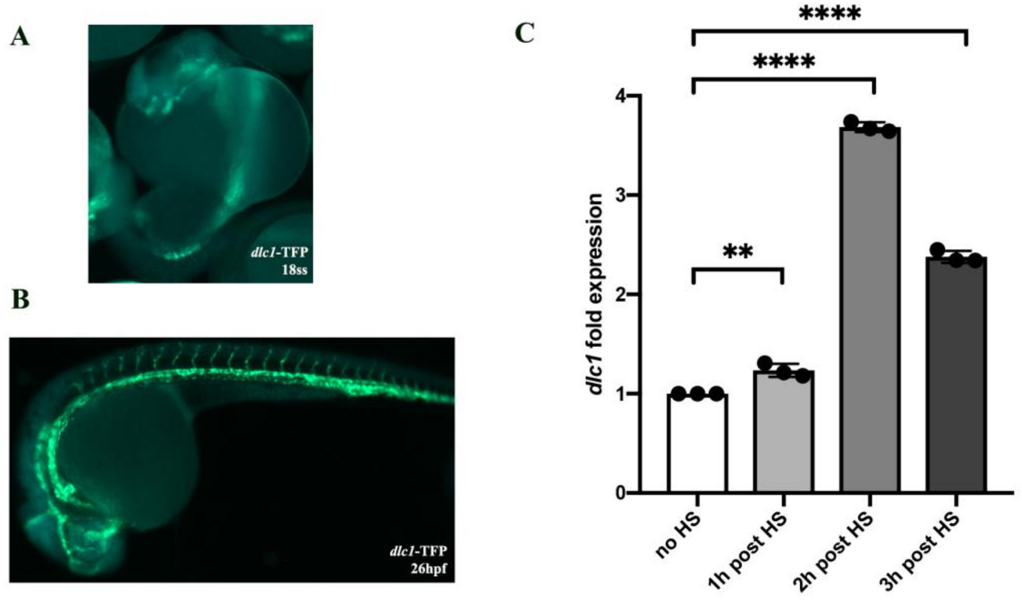

**Suppl. Fig. 2 Validation of the *hsp-dlc1* line**

A) and B) show heat shocked embryos at indicated timepoints. *dlc1*-specific TFP fluorescence (blue) could be visually detected as early as 18 ss (2.5h after heat shock) in the whole embryo (no tissue-specificity). In both pictures, the vasculature is labelled by *kdrl:eGFP*. C) shows the relative *dlc1* expression (fold change) compared to non-heat shocked control embryos over the timecourse of 3h after the heat shock. Each timepoint consists of a pool of 10 embryos per datapoint (n = 30). Data represents the mean ±SEM. Statistical significance was determined using a 1-way ANOVA with Dunnett's post hoc correction. **p < 0.01; ****p < 0.000.1

**Supplemental Fig. 3**

**A**

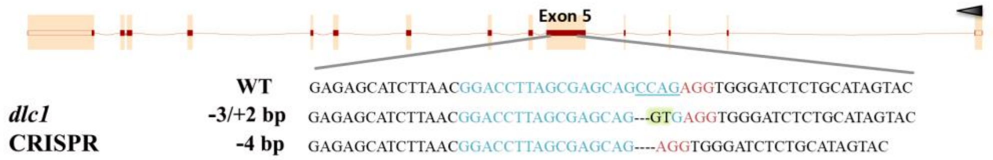

**B**

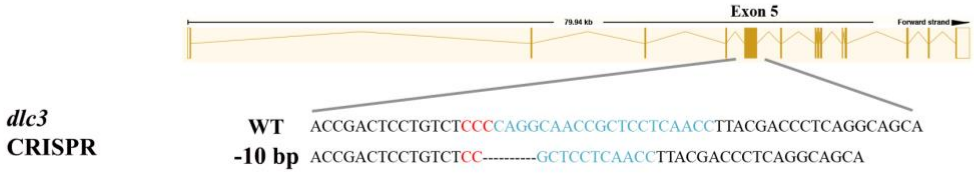

**C**

***dlc1* and *dlc3* MO strategy**

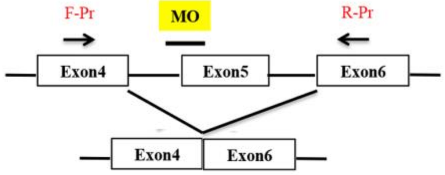

| Target gene | WT product | Morpholino PCR product |
|---|---|---|
| *dlc1* | 1811 bp | 234 bp |
| *dlc3* | 1534 bp | 227 bp |

**D**

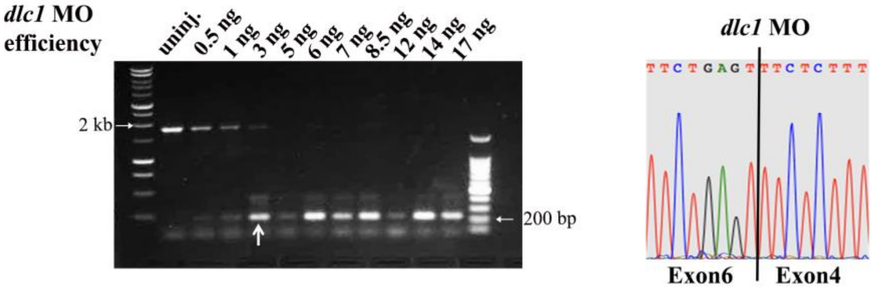

**E**

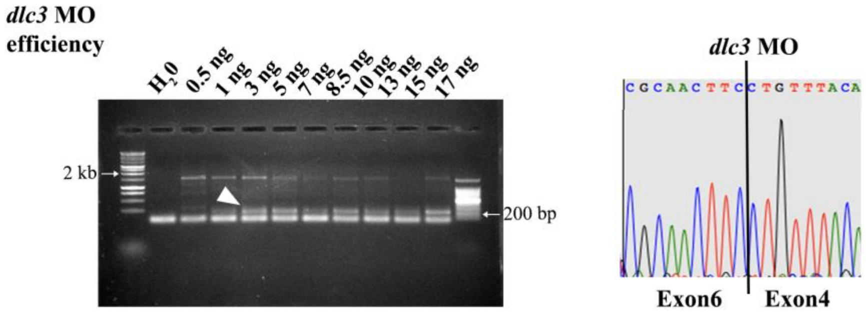

**Suppl. Fig. 3 CRISPR/Cas9-mediated knockout (KO) and Morpholino (MO) design**

A) shows the CRISPR/Cas9-mediated KO strategy for *dlc1* in Exon5. The nucleotide sequence around the target site (highlighted in turquoise with the PAM motif in red) in the WT *dlc1* gene and the two mutated sequences (-3/+2bp and -4bp) that both generated a premature stop within the same Exon5 (after 174 translated AA) are shown. B) shows the CRISPR/Cas9-mediated KO strategy for *dlc3* in Exon5. The nucleotide sequence around the target site (highlighted in turquoise with the PAM motif in red) in the WT *dlc3* gene and the mutated sequence (-10bp) that generated a premature stop within the same Exon5 (after 163 translated AA) are shown. C) shows the splice blocking MO strategy between Intron4 and Exon5. The locations of the primers that were used to amplify the affected region are indicated in red and the expected PCR products for the WT or splice blocking MO are listed in the adjacent table. The *dlc1* and the *dlc3* MOs targeted a similar region compared to the affected one by CRISPR/Cas9 mutagenesis. The efficiency of the MOs is shown using Gelelectrophoresis. The loss of Exon5 is shown by the appearance of a smaller gene product (indicated by a white arrow for the *dlc1* MO and a white arrowhead for the *dlc3* MO). Corresponding histograms for the sequenced small bands show the seamless transition from Exon4 to Exon6 and the loss of Exon5 (the reverse Primers were used for sequencing).

**Supplemental Fig. 4**

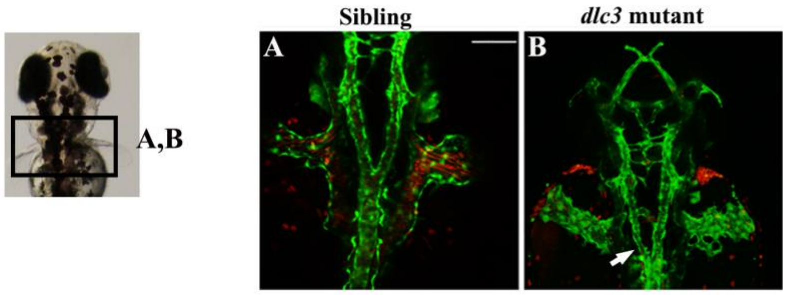

**Suppl. Fig. 4 In rare cases, the *dlc3* mutant shows a vascular phenotype with impaired LDA formation**

A) and B) show dorsal views of the LDA in WT siblings (A) and *dlc3* mutants (B) on a *kdrl:*eGFP transgenic background. The mild stenosis of the LDA in the *dlc3* mutant is marked by the white arrow. The presence or absence of bloodflow in the *dlc3* mutant (B) compared to the WT sibling (A) throughout the LDA and the CCVs are demonstrated by the *gata1*:DsRed transgene.

**Supplemental Fig. 5**

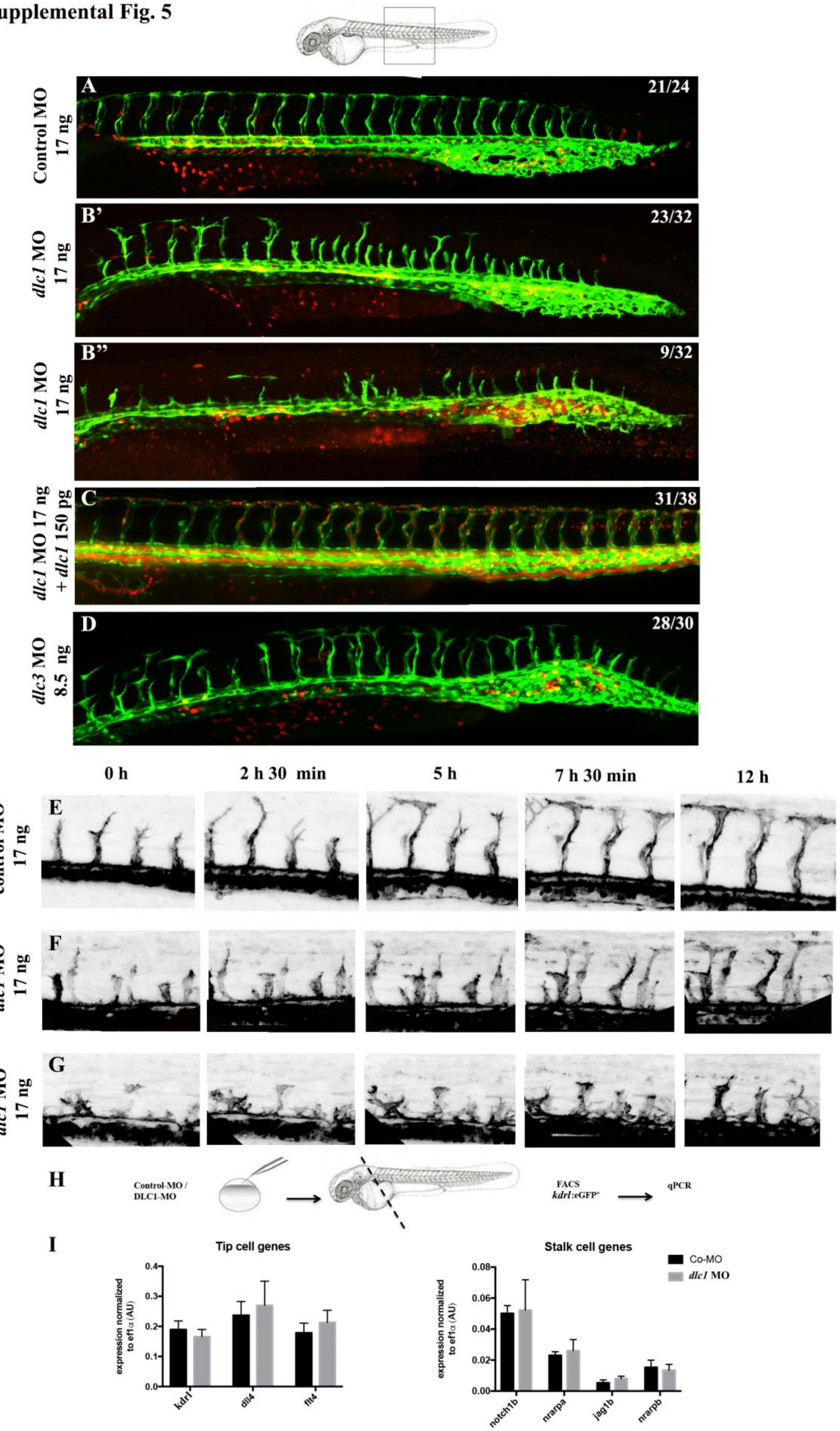

**Suppl. Fig. 5 Transient knockdown of *dlc1* and *dlc3* greatly impairs ISV growth**

A), B) and D) show lateral views of the ISV in control MO injected embryos (A)), *dlc1* MO injected embryos (B and B')) and *dlc3* MO injected embryos (D)) on the *kdrl:eGFP/gata1:dsRed* background at 33 hpf. C) shows the rescue of the *dlc1* MO induced ISV phenotype by co-injection with full length *dlc1* mRNA. E) – G) show sequential images from selected timepoints of timelapse movies (Supplemental Movies 1-3) using *fli1a:lifeact*-GFP embryos. E) shows the ISV growth of control MO injected embryos and F) and G) the ISV growth of *dlc1* MO injected embryos. The trunk and tail of control and *dlc1* MO injected embryos were dissected and ECs sorted (H). qPCR analysis (biological triplicates, N = 3) of tip and stalk cell identity genes is shown in I). qPCR data was analyzed using an unpaired, 2-tailed t test for each individual comparison. P values: *kdrl* p = 0.332 (n.s); *dll4* p = 0.571 (n.s); *flt4* p = 0.308 (n.s); *notch1b* p = 0.863 (n.s); *nrarpa* p = 0.529 (n.s); *jag1b* p = 0.137 (n.s); *nrarpb* p = 0.631 (n.s)

**Supplemental Movies**

**Suppl. Movie 1:** Time-lapse movie of Tg(fli1a:lifeact-GFP)$^{mu240}$ embryos (24 ss onwards) injected with a standard control MO (17 ng) showing normal ISV development. A 2.5µm z-stack (total imaged depth 80 µm) was obtained with a 20x objective every 10min for 13h during which the embryos were maintained in a heated chamber at 30°C. The z stacks were combined to a maximum projection movie using the FIJI software. Supplemental Figure 5E shows single images at selected timepoints.

**Suppl. Movie 2:** Time-lapse movie of Tg(fli1a:lifeact-GFP)$^{mu240}$ *dlc1* morphant (17 ng) showing abnormal ISV growth from 24 ss onwards. Excess short sprouts as well as a stalling growth behavior are shown. A 2.5µm z-stack (total imaged depth 80 µm) was obtained with a 20x objective every 10min for 13h during which the embryos were maintained in a heated chamber at 30°C. The z stacks were combined to a maximum projection movie using the FIJI software. Supplemental Figure 5F shows single images at selected timepoints.

**Suppl. Movie 3:** Time-lapse movie of a second Tg(fli1a:lifeact-GFP)$^{mu240}$ *dlc1* morphant (17 ng) showing abnormal ISV growth from 24 ss onwards. Sprouts form vascular plexi and fail to elongate. A 2.5µm z-stack (total imaged depth 80 µm) was obtained with a 20x objective every 10min for 13h during which the embryos were maintained in a heated chamber at 30°C. The z stacks were combined to a maximum projection movie using the FIJI software. Supplemental Figure 5G shows single images at selected timepoints.

**Suppl. Movie 4:** Brightfield movie showing the heartbeat of *a dlc1* mutant embryo at 2.5 dpf. The blood is visible in the brightfield channel and is pumped back and forth from the atrium into the ventricle without leaving the heart. The movie was taken of a mutant embryo in 1xE3 anesthesized with a low dose of Tricaine and a 10x objective.

**Suppl. Movie 5:** Shows the same embryo as in Suppl. movie 4 visualizing *kdrl:eGFP* expression. The movie shows that the heart and vasculature are physically connected, but contain a visible obstruction in the region of the OFT. The movie was taken of a mutant embryo in 1xE3 anesthesized with a low dose of Tricaine and a 10x objective.

### Supplemental Tables

**Suppl. Table 1**: Truncated mRNA constructs for *dlc1*, the contained active domains and their effect after injection

| **injected truncated *dlc1*** | **active domains** | **effect (doses injected)** |
|---|---|---|
| **ΔFAT-GAP-START** | SAM (AA 1-70) | no effect (50 pg - 250 pg) |
| **ΔGAP-START** | SAM and FAT<br>(AA 1-648) | no effect (50 pg - 250 pg) |
| **ΔSAM** | FAT-GAP-START (AA 70-1128) | lethal (100 pg and 200 pg) |
| **ΔSAM-FAT** | GAP-START<br>(AA 648-1128) | mild ISV phenotype<br>CCV phenotype (200 pg) |
| **ΔSTART** | SAM-FAT-GAP<br>(AA 1-894) | no effect (200 pg) |
| **ΔFAT-GAP-START + ΔSAM** | all domains | lethal |
| **ΔGAP-START + ΔSAM-FAT** | all domains | lethal |
| **ΔFAT-GAP-START + ΔSAM-FAT (→□FAT)** | SAM-GAP-START | no effect (100 pg and 200 pg each) |

**Suppl. Table 2**: Primers used for ISH, qPCR and genotyping

| | Forward primer | Reverse primer |
|---|---|---|
| For ISH | | |
| dlc1 | TCAAATTGAGGCGAAGGAGGC | GCTCTTGACCGTGTGCTTTTT |
| dlc2 (stard13b) | TCCAGATTTCCAACCGCCTC | GCCTCCCTGTGGCATACTTT |
| dlc3 (stard8) | TTGAGGAATGGTCACGCTCC | TGGATGCTTTGAGGCAGAGG |
| for qPCR | | |
| bmp4 | CCGTGAGAGGATTCCATCAT | TGTTTATCCGATGCAAACCA |
| dlc1 | GGAGCCTGTTCTCATCAGCC | TGACAGCACTTTGACATAGGATTGC |
| dlc3 | GAGTTGGCGAAGGAGTTTGG | TTGCCTCCAGCTCCGC |
| dll4 | AAACCTGGAGAGTGTGTATGC | TGGTCACAGAACAGTCCACC |
| ef1a | GAGAAGTTCGAGAAGGAAGC | CGTAGTATTTGCTGGTCTCG |
| kdrl | CTCCTGTACAGCAAGGAATG | ATCTTTGGGCACCTTATAGC |
| flt4 | AGGCTGAAGGAGTGTCAAGG | GAATGGATGCTCTGGGTCTCG |
| jag1b | AGACCGCCAGTGAAAGTGG | GCCGTAGTAGTGTTCAAGGC |
| myl7 | GCCCATAAACTTCACTGTCTTCC | GACAACTCCTGTGGCATTAGGG |
| notch1b | AGGCGTCTTCCAGATTTTGA | CCTCGACCGCCAGTCTT |
| nrarpa | AACATGACCAACTGCGAGTTTAAC | GAATATCGGCTCCAAATTTAACCA |
| nrarpb | CCGCAGAGGTATCGGAGCG | CTGAAGGAGGGAAATAAGCCAGG |
| notch1b | AGGCGTCTTCCAGATTTTGA | CCTCGACCGCCAGTCTT |
| plxnd1 | TGAAGAACGTGGAGCATGGG | TCGATGTATGTGGCGTCTCC |
| for genotyping | | |
| TFP | GGCGTAATCAAGCCCGACAT | TGGTCTTGAAGTCAACGCGG |
| dlc1 mutant | TGGTGTGCATGTGACTGTGCTGTG | GCTCTTTGCTCTTGACCGTGTGCT |
| | sequencing: GTGCCATAAGTGGCCGTTGGAC | |
| dlc3 mutant | GAGGCTTGGGAATGGGTTCA | TCATTCTCACTGTCCCGCTTC |
| MO efficiency | | |
| dlc1 | ACCCTAAACAAGTGCGCTCT | CAGCTGAAGCCATGTTTGTT |
| dlc3 | CCCTGAACAAATGTGCTGCG | CAGTTCCAGCCATGCTTGCT |
